# Continuous Rate Modelling of bacterial stochastic size dynamics

**DOI:** 10.1101/2020.09.29.319251

**Authors:** Cesar Nieto, Cesar Vargas-Garcia, Juan Pedraza

## Abstract

Bacterial division is an inherently stochastic process. However, theoretical tools to simulate and study the stochastic transient dynamics of cell-size are scarce. Here, we present a general theoretical approach based on the Chapman-Kolmogorov formalism to describe these stochastic dynamics including continuous growth and division events as jump processes. Using this approach, we analyze the effect of different sources of noise on the dynamics of the size distribution. Oscillations in the distribution central moments were found as consequence of the discrete translation invariance of the system with period of one doubling time, these oscillations are found in both the central moments of the size distribution and the auto-correlation function and do not disappear including stochasticity on division times or size heterogeneity on the population but only after include noise in either growth rate or septum position.

## Introduction

Recent experiments involving time-lapse microscopy [1], single-cell tracking [2, 3], and gene tagging [4] have revealed how cell size stochasticity, and division events play an important role in the random fluctuations of bio-molecular concentrations [5–8]. This with important consequences for phenotype variability and cell heterogeneity over a clonal population of microorganisms [9].

Methods approaching the bacterial size control include the discrete stochastic maps (DSMs) [10]. These models define the known as *division strategy* as a map that takes cell size at birth *s_b_* to a targeted cell size at division *s_d_* trough a deterministic function *s_d_* = *f* (*s_b_*) plus stochastic fluctuations that have to be fitted from experiment. DSMs, however, are unable to reproduce cell size transient dynamics at arbitrary infinitesimal time intervals without further extension.

To solve this continuous dynamics of the distributions describing stochastic processes usually the Chapman-Kolgomorov formalism (CK) is used [11]. CK solutions corresponds to the distributions of all the possible stochastic hybrid trajectories at a given time. Among the processes that can be modeled by CK it can be included continuous size growth and division as a jump process requiring the definition of a continuous rate of division. In fact, these models are also known as Continuous Rate Models (CRM) [10]. Recently, [12] proposed a power-law function to explain observations in E. coli bacteria cells. Instead, [13] suggested a convoluted function of size and cycle progression is required. [14] proposed a deconvoluted version by introducing division as a multistep process where the occurrence rate of these steps is a function of the size. Despite these recent attempts, there is a lack of a complete formalism describing the phenomena related to bacterial division.

Here, we propose the CK formalism as a framework for studying single-cell transient dynamics. We present how to overcome the non-locality of the division jumps and how to model the division steps. These steps being analogous to the experimental accumulation of FtsZ to trigger the division [15, 16]. These equations are solved using both simulations and numerical methods. Analytical expressions are also presented in the cases where it was possible. We also present modifications to the division rates to define multiple division strategies. Then, we will show how stochasticity on division influence the size distribution dynamics and how this dynamics changes considering additional sources of noise like a distribution of initial sizes with finite variance, the noise in septum position and in cell-to-cell growth rate. We discuss how this approach could be coupled to simulations of gene expression.

## Theoretical details

### The population balance Equation

Consider the distribution *p*(*s*; *t*) of sizes *s* at a given time *t* solving the Chapman-Kolmogorov equations (CKE). This distribution is related to the histograms of bacterial populations. Some attempts have described the dynamics of this distributions including effects due to the increasing of population number and the mother-daughter correlations [17–19]. In our framework, we consider a cell population with an asymptotically large number. After each division, only one of the descendants is tracked such as the number of bacteria in the population remains constant in similar way observed in experiments such as those using *Mother Machine* micro-fluids [3]. Then, *ρ*(*s*; *t*) can be normalized:

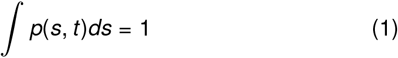

To describe the distribution dynamics, let an individual cell grow in size *s* as per

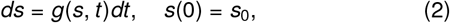

where *g*(*s*, *t*) is the size change per unit time. Some studies have considered the constant growth [17, 20, 21] we will focus con exponential growth rate (*g*(*s*) = *μs*) with *μ* being the growth rate and *s*_0_ is the initial cell size. Since (2) defines a deterministic process, the change in the size distribution *p*(*s*, *t*|*s*_0_) conditioned on initial size *s*_0_ can be obtained by solving

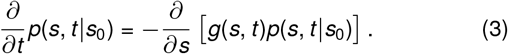

Expression (3) is known as the CKE in its differential version (dCKE), and its solution for this deterministic process is given by

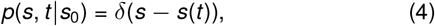

where *s*(*t*) is the solution of equation (2).

Considering division as a jump process switching, at a time *t*, from cell size *s′* to size *s* at rates *W* (*s|s′*, *t*), equation (3) can be written as

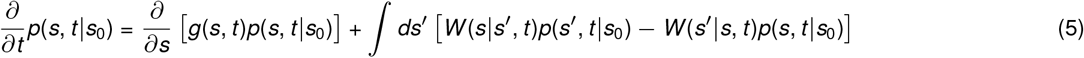

If perfect symmetric splitting is considered through the condition (*δ*(*s′* − 2*s*)) and |*W*(*s′|s*,*t*)| = *h*(*s*, *s′*), the division rates *W* (*s|s′*, *t*) can be written as:

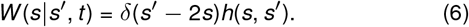

Some studies have explored the particular case where *h*(*s*) = *ks* with *k* being a constant and discarding the dependence upon *s′* [17, 22]. This, resulting on:

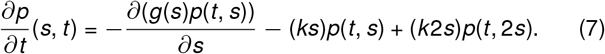

Which is also known as the Population Balance Equation for a fixed population number [17]. Some theoretical methods like moment closure were used in past studies to solve (7) but stable solutions were not found already [17].

### The Chapman-Kolmogorov equation including division events

To overcome these instabilities, other studies reparametrize the size distribution adding an additional variable: the number of divisions *n* ∈ {0, 1, 2, …} [22]. In this case, the probability distribution is now *p*(*s*, *n*; *t*) with transition rates satisfying:

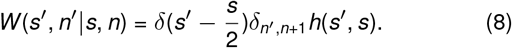

This is, after division, not only the size is halved but the number of divisions *n*, increase by one unit. In the particular case where *h*(*s*, *s′*) = *ks*, the associated CKE is:

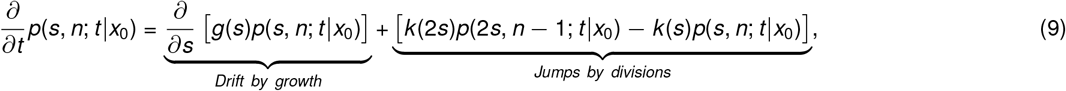

where *x*_0_ = (*s*_0_, *n*_0_).

The inclusion of the new variable *n* breaks the non-locality of the operator *W* (*s|s′*, *t*) that makes (5) hard to solve. Instead *W*(*s*, *n|s′*, *n* − 1, *t*) performs jumps between independent sub-spaces that can be merged together later with marginal sums.

Using this new variable *n*, exponential growth and *s*(0) = *s*_0_ equation (9) has closed solutions

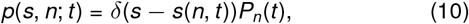

where *P_n_*(*t*) is the probability to get divided *n* times at time *t*. We will present its associated equation later. *s*(*n*, *t*) corresponds to the bacterial size after *n* divisions at a time *t*. To explain how to solve this size, let us consider the size *s*(1, *t*) at time *t* after one division. If *t*_1_ < *t* is the time when division occurs, this size is:

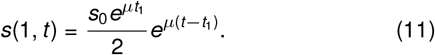

So, having the general sequence of division times 0 < *t*_1_ < *t*_2_ *<* … < *t*_*n*−1_ < *t_n_* < *t*, *s*(*n*, *t*) satisfies:

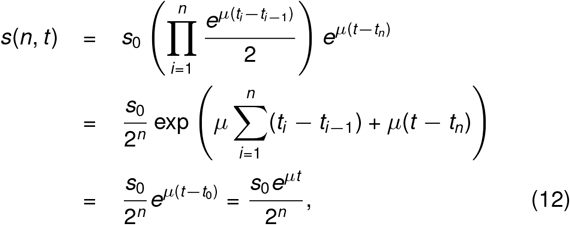

where we used *t*_0_ = 0 and the telescopic properties of the sum.

Using this result (12) and (10), (9) is separated to the system of equations:

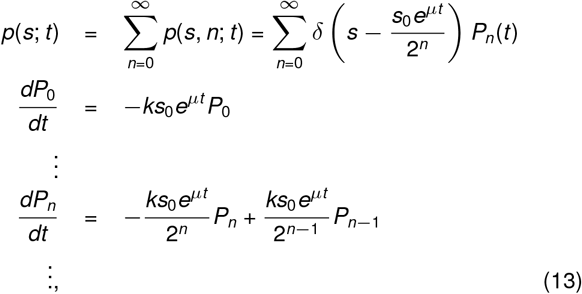

defining the dynamics of every *P_n_*(*t*).

### The division strategy

Focusing on the jump process between the state *n* − 1 to *n*, or simply, one division, we can modify the space *n* ∈ 0, 1, 2, … to *n* ∈ {0, 1} and truncate (13) to:

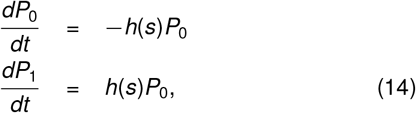

Where a general size dependent splitting rate function *h*(*s*) can be used. This system can be integrated under the initial conditions *P*_0_(0) = 1 and *P*_1_(0) = 0. Thus, the probability *P*_1_(*t*) ‒to simplify notation, *P*(*t*)–that the cell divides in the time interval (0, *t*) evolves according to:

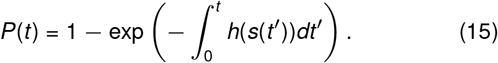

In the particular case of *h*(*s*), proportional to the size, this is, *h*(*s*) = *ks*, assuming exponential growth as well, the integration on time has to be done using the implicit formula:

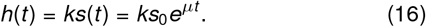

Once *P*(*t*) is obtained, the probability density function *ρ*(*t*) for the time of division can be obtained as:

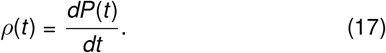

A transformation of variables, allows us to get the distribution of sizes at division *ρ*(*s_d_*):

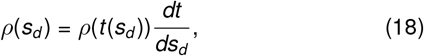

where, if we assume exponential growth 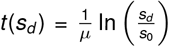, then, 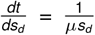. Using this *ρ*(*s_d_*), one can calculate the mean size at division by integrating:

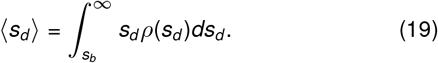

Hence, we can calculate the mean added size per cell cycle 〈Δ〉 = 〈*s_d_*〉 − *s_b_* as a function of the size at birth *s_b_*. This relationship defines the division strategy.

### The multi-step Single division

In the general case, division does not correspond to a single jump process, instead, division occurs once bacteria have reached some goal steps *M*. If the occurrence rate of these steps is proportional to the cell size *s* by the constant *k_d_*, the probability *P_m_*(*t*) of having done *m < M* steps at time *t* can be modelled following:

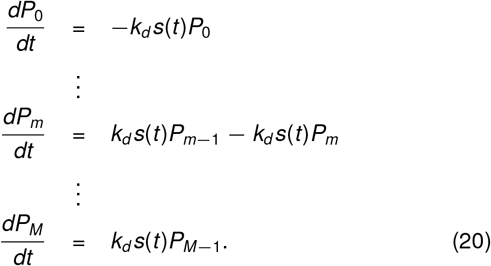

*P_M_* is the probability of reaching the target steps *M* or equivalently, the probability of a division event to occur. Once the division event happens, the process is reset to zero steps and size is halved.

Using this *P_M_*(*t*) and the growth regime (4), if the procedure (18) is followed, the probability density *ρ*(*s_d_|s_b_*) of size at division *s_d_* given the size at birth *s_b_* in a cell cycle, satisfies:

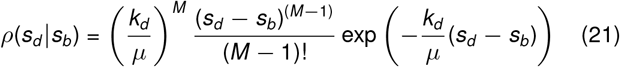

Defining the added size before division Δ = *s_d_* − *s_b_*, from, we observe that 〈Δ〉 = 〈*s_d_*〉 − *s_b_* is independent on the size at birth *s_b_* and is related to the growth rate *μ*, the objective steps *M* and the step occurrence rate *k_d_* per size unit, satisfying:

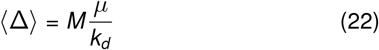

### The solution of the CKE including multiple divisions

Using a similar procedure as (13) now with the additional variable *m*, the probability of have a size *s*, have done *m* division steps and *n* divisions up to time *t* can be written now as:

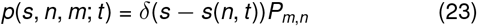

Cell size *s*(*n*, *t*) follows, again, the equation (12). While, the probability *P_m,n_*(*t*) of have done *m* division steps and *n* division events up to time *t*, can be estimated through the master equation system:

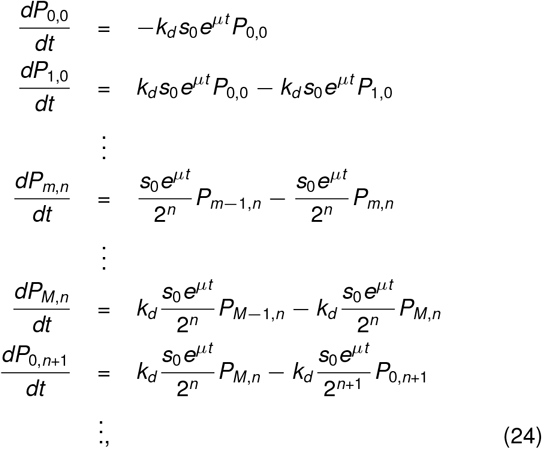

where we showed the selection rule defining the divisions as jumps between states (*M*, *n* − 1) to (0, *n*) and the division steps as jumps between states (*m*, *n*) to (*m* + 1, *n*). These jumps happening at rate *h* = *k_d_ s*(*n*, *t*) with *s*(*n*, *t*) following the equation (12).

## Numerical estimation of size dynamics

Solution of (24) can be obtained using different methods. Analytically, one can start from the initial conditions:

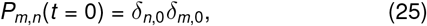

with *δ*_*i,j*_, the Kronecker delta. Hence, *P*_*m*,*n*_(*t*) can be obtained knowing *P*_*m−*1,*n*_(*t*) with *m* ∈ 1, …, *M* and *P*_0,*n*_(*t*), can be estimated from *P*_*M*,*n*−1_(*t*). Both, using, using the closed recurrence expression:

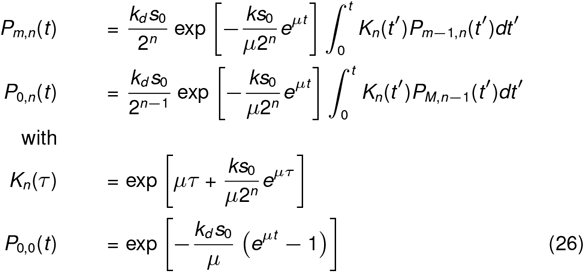

### The Finite State Projection Algorithm

In general, in (24), while the number of possible steps *m* ∈ {0, 1, …, *M*} has finite cardinality, the number of possible divisions *n* ∈ {0, 1, …} is infinite. Thus, making impossible the complete solution of (24) using methods like Matrix exponential. As we explained before [22], this infinite set can be projected into a finite set using the known Finite State Projection (FSP) algorithm [23]. Using this approach, the number of equations in (24) are truncated up to a maximum divisions *N* and the number of possible division states is now finite. Hence, known methods for solving these finite systems can be used to estimate size dynamics during infinitesimal periods of time.

From *P_m,n_*(*t*), the size distribution *ρ*(*s|s*_0_) given the starting size *s*_0_ and the moments of this size distribution: the mean size 〈*s*〉 and the variance var(*s*) = 〈*s*^2^〉 − 〈*s*〉^2^, can be estimated from the equations:

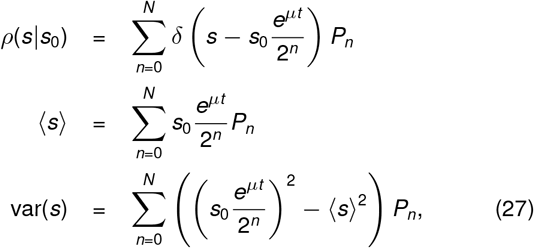

with 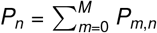 and *δ*(*x*), the Dirac-delta distribution.

The computing of these moments was done considered that all cells began at initial size *s*(0) = *s*_0_, this is, *ρ*(*s*_0_) = *δ*(*s*_0_ − *s*(0)). However, if a general density function *ρ*(*s*_0_) is considered, the size distribution *ρ*(*s*) is a convolution of solutions of (27):

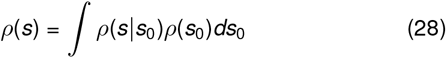

## Stochastic simulation of size dynamics

Let the single step process shown in (14). While (14) was presented to modeling the division as a single step process, in general, these equations are also valid for a division step. Setting explicitly the dependence *h* = *k_d_s*, the system describing the single step process is now:

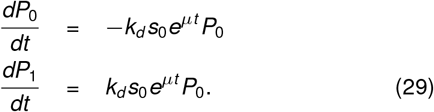

If *P*_0_(0) = 1 and *P*_1_(0) = 0, *P*_1_(*t*), or simply *P*(*t*), has solution:

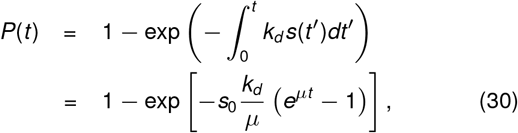

while the associated density function is:

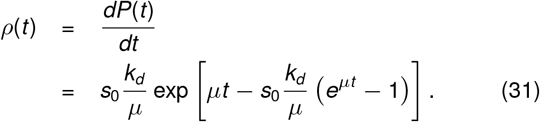

The main idea behind the stochastic simulation algorithm is to generate random time events *τ_s_* distributed as (31). Following the Gillespie’s method [24], we generate a random number *r* uniformly distributed in the interval (0, 1) and from the cumulative function (30), *τ_s_* is obtained matching *P*(*t*) and *r* thus, solving for *t* :

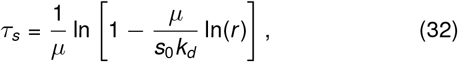

where we take advantage of the fact that 1 − *r* is distributed as *r*. This *τ_s_* is the time to the occurrence of the next division step.

## Additional details to model the cell division

### The asymmetric splitting

The main assumption proposing (12) is that after each division, the cell size is perfectly halved. However, this is not the case in a realistic situation. Experimentally, some stochastic fluctuations in the septum position are found. In some growth conditions, this noise can be as high as 5% [25, 26].

Considering again the size at time *t* after one division, if the division occurred at time *t*_1_ < *t*. If the size is not perfectly halved but multiplied by a random variable *b*_1_, centered on 0.5 and also known as division ratio, the size at time *t* is now:

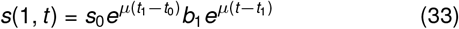

If the sequence of division ratios {*b*_1_, *b*_2_, …, *b_n_*} is known, the size after *n* division at time *t* is given by:

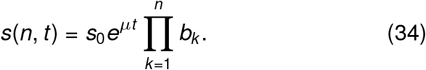

Theoretically, *b_k_* can be approximated to a beta distributed variable centered on 0.5 with variance fitted from experiments.

### Cell-to-cell noise in growth rate

Other important stochastic variable is the cell-to-cell growth rate [27, 28]. This fluctuation can be as high as 10% [3]. We can assume that after a division *k*, a new growth rate *μ_k_* is chosen randomly from a distribution centered on *μ* and cell grows during that cycle with rate *μ_k_*. If symmetric splitting is considered again, the size at time *t* after *n* divisions is now:

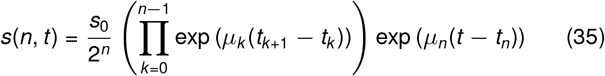

Numerically, we modeled these *μ_k_* as a gamma distributed variable centered on the mean growth rate *μ*, with the experimental variance and no correlation with past cycles. This last assumption might be incorrect and these correlation between cycles can be studied in deeper studies [26].

### A general CKE incluing additional souces of noise

A general CKE, assuming exponential growth, can be modeled with growth rate distributed with given distribution *ρ*(*μ*):

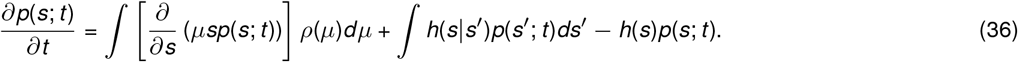

The division rate *h*(*s|s′*) in general depends on hidden variables like the number steps but can estimated, at least numerically [29–31]. This In the case of single step division, this rate can be written as *h*(*s|s′*) = *ksρ*(*s*, *s′*) with ∫ *ρ*(*s*, *s′*)*ds′* = *ρ*(*s*) and *ρ*(*s*) being the distribution of size at birth. In the therm *ρ*(*s*, *s′*) non-symmetric division can also considered. *h*(*s*), in the last therm, is obtained from *h*(*s*) = ∫ *h*(*s′|s*)*ds′*. We already do not have a closed solution to (36) and numerical methods could be hard to implement.

### Different division strategies

Depending on the mapping *s_d_* = *f*(*s_b_*), or traditionally, between the relationship between added size Δ = *s_d_ − s_b_* and *s_b_*, three main division strategies have been defined for exponentially growing bacteria: the *timer*, *adder* and *sizer* strategies [32]. Differing from the slope of Δ vs *s_b_* for timer this slope is +1; −1 in sizer strategy and for null-valued for adder.

The adder strategy, observed for instance, in *E. coli* and *B. subtilis* [3], is considered the most common strategy in bacteria. In some bacterial populations, however, division strategies with intermediate slopes for Δ vs. *s_b_* have also been observed [14, 32]. This has led to the definition of the *timer-like* strategy, for slopes between 0 and 1, and of the *sizer-like* strategy, for slopes between −1 and 0.

These deviations from the adder can be obtained if the step rate (*h*) is not proportional to the size but proportional to a power (λ) of the size [14]. This is

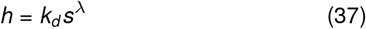

A multi-step process similar to that described by (24) can be proposed. As was explained in past studies [14], if division is triggered by the occurrence of *M* steps happening at rate (37), the distribution of size at division *s_d_* given the size at birth *s_b_* is [14]:

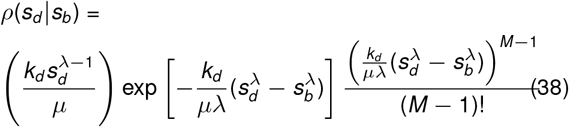

Timer strategy (Added size has a slope of −1 with size at birth) is obtained if λ → 0 and sizer strategy (Added size is proportional to the size at birth) is obtained if λ → ∞. The adder is obtained when λ = 1.

Considering the non-linear step rate given by (37) and a similar method like (31), the general stochastic time is given by:

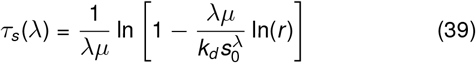

### The mean cell size at birth

The main variables defining the mean cell size are the growth rate *μ*, the number of division steps *M* and the division steps occurrence rate *k_d_*. If adder strategy is considered (λ = 1), the mean added size 〈Δ〉 follows the relationship (22).

Since there is already a discussion about the nature of *k_d_*, the inference of its actual value is not straightforward. For the adder, this *k_d_* can be inferred from their mean added size using (22) and by observing that this 〈Δ〉 is independent on the size at birth *s_b_*. In different division strategies (λ ≠ 1), 〈Δ〉 is now function of *s_b_*. Now, the typical size as explained in past studies [14, 17], 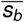, is the size at birth that is perfectly doubled by the division strategy:

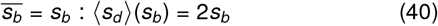

This, since after division, the cell, with 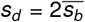, splits on a half and the size of its offspring is 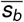 again.

In general, 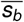, is dependent on *k_d_*, *μ* and λ and is one of the variables that can be measured most easily if we assume that this 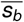 is actually the mean size at birth in a steady growing cell population. Hence, other variables like *k_d_* can be estimated from this 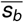 using (40) and root-finding algorithms.

## Illustrating examples

To observe the dynamics of the probability of get divided *n* times at time *t*, we present, in Figure 1, time trends of some *P_n_*(*t*)s for different λs and *M*s with initial condition *P_n,m_*(0) = *δ_n_*_,0_*δ_m_*_,0_. .

**Figure 1:**
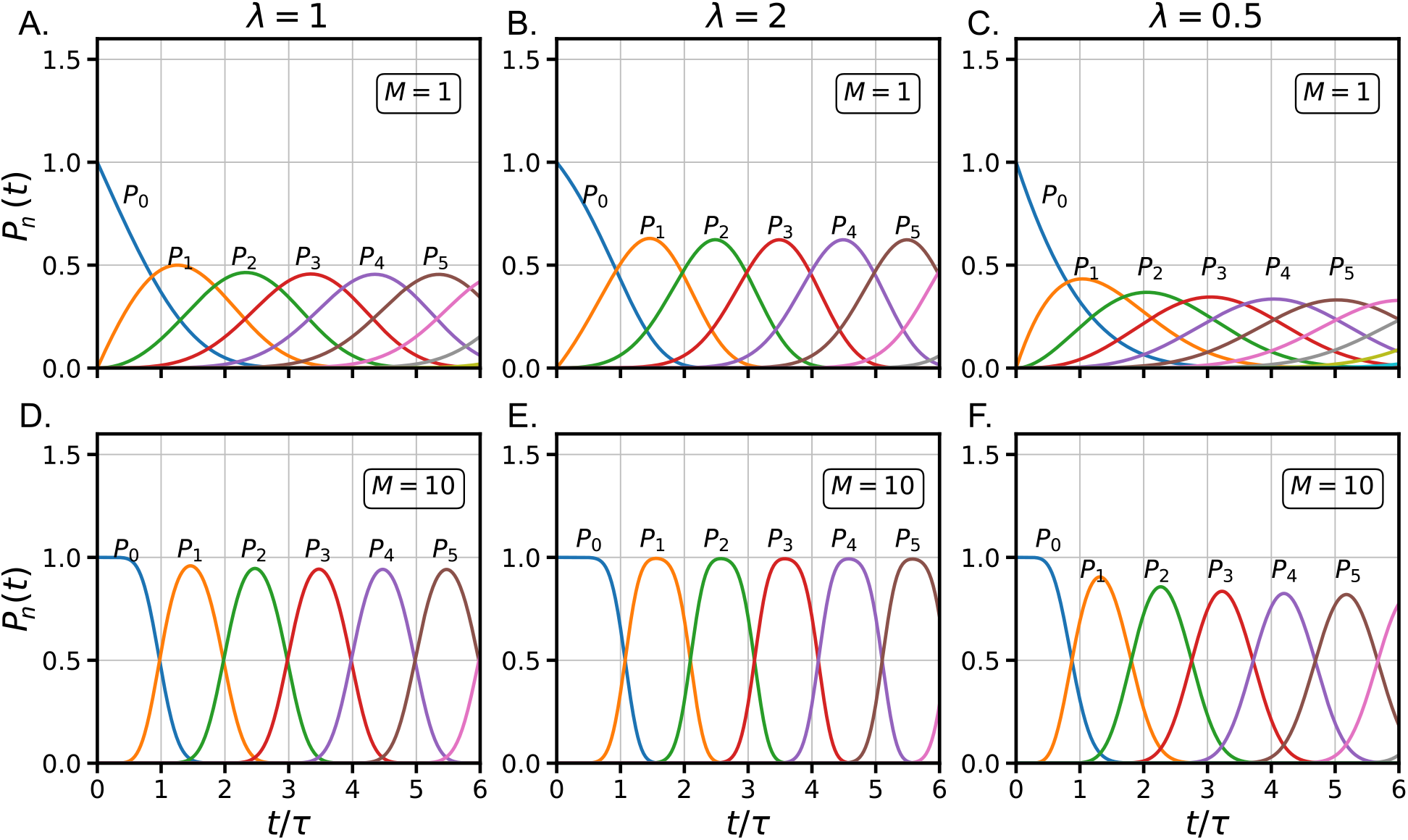
Time dynamics of the first *P_n_* for different division strategies and division steps (*M*) with 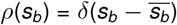.

A numerical analysis of the behavior in Figure 1 allows us to find that, in the limit of *t* → ∞, the distribution of *P_n_*s satisfies

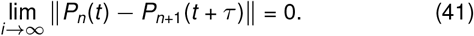

Which implies asymptotic invariance of the system under translation on, simultaneously, *n* → *n* +1 and *t* → *t* + *τ* over the *P_n_*. Since 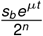 also satisfies this invariance, we expect *ρ*(*s|t*, *s*_*b*_) to show periodic properties in the limit *t* → ∞. This periodicity was already discussed in some theoretical papers [33–35]. This convergence is shown in Appendix A.

Three different scenarios in size dynamics can be explored using our formalism. Figure 2 a. shows the mean size dynamics obtained from a simulation of 5000 cells, all of them with the same starting size and beginning with zero division steps or equivalently, starting from their most recent division. We assumed that they have the same growth rate and get split perfectly symmetric. Simulation was done using the stochastic times (32) and numerical estimation was done solving the master equation (24, both of them, having perfect adjustment to each other. Ten examples of cell cycles were plotted on the background to observe how variable the distributions are. The dynamics of this variability, quantified by the coefficient of variation 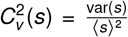, is also shown in Figure 2.e. As main effect observed, we can highlight the oscillations in both 〈*s*〉 and 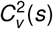 with period equal to *τ*, the doubling time. The oscillations in the 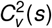 present their peaks just when bacteria are dividing on average and their valleys when bacteria are growing. The dynamics of the size distribution can be seen in supplemental video 1.

**Figure 2:**
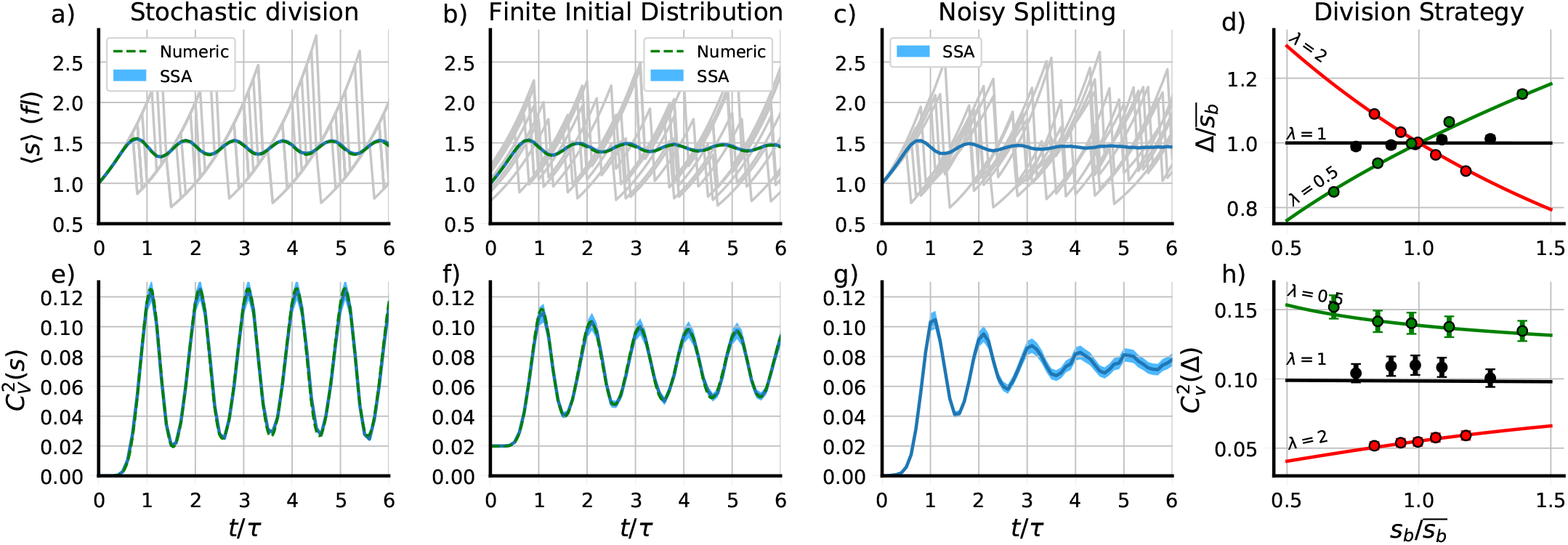
Main properties of bacterial cell division explored using PyEcoLib. a) Mean cell size ⟨*s*⟩ and e) Its variability 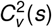 along the time considering only stochastic division. b) Mean cell size and f) Its variability along the time considering both stochastic division and an initial size distribution with finite variance. c) Mean cell size and g) Its variability along the time considering stochastic division and noise in cell-to-cell growth rate and septal position. d) mean added size Δ vs the size at birth *s_b_* and h) the fluctuations 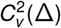 vs *s_b_* for different division strategies. Timer-like (λ = 0.5), adder (λ = 1) and sizer-like (λ = 2). Simulations (dots) and numerical estimations (lines) are shown. *M* = 10 division steps were considered in all cases.

In Figure 2. b., the mean dynamics corresponds to cells with an initial distribution with finite variance 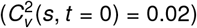. This distribution was assumed to be Gamma distribution since it is well defined from its mean size and the 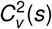. Simulations, on the other hand, were modified by using random initial sizes, numerical estimation was done by performing the convolution (28) and approaching the integral to a numerical Riemann sum. Dynamics on cell size variability are also presented in Figure 2. f. Similar oscillations were found in both the mean and the variability but with less amplitude than the first case. The dynamics of this distribution appears in supplemental video 2.

A third scenario consists on the assumption that bacteria do not split perfectly on a half but in a beta distributed independent stochastic variable centered on 0.5 and with a given variability 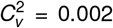. Growth rate could be considered stochastic as well with variability set to 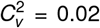. The dynamics of the mean size (Fig 1.c.) and its variability (Fig 1.g.) are presented. We plotted ten cycles in the background of Figure 2.c. to show some examples of typical single-cell size dynamics. Since this noise is not considered in equation (24), the numerical approach is hard to compute and thus not presented in Figure 2. The distribution dynamics can be seen in supplementary video 3.

We also estimate the division strategy using both simulations and numerical estimations. Data from stochastic simulations can be obtained using the stochastic division times (39) and exponential growth (4) while the trends in added size and its variability can be obtained from the distribution of size at division (38) being both dependent on the exponent λ. In Figure 2 d. we present the mean added size *δ* as function of the mean size at birth *s_b_* for three different λ. These λ were chosen to represent three of the most important division strategies: timer-like (0 < λ < 1, where we choose λ = 0.5) with its characteristic positive slope on Δ vs *s*)*b*, Adder (λ = 1) with no correlation between Δ and *s_b_* and sizer-like (1 < λ < ∞, where we choose λ = 2) with a negative slope in Δ vs *s_b_*. Fluctuations over these trends are also shown in Figure 2 h. where it can be seen that sizer-like shows positive correlation in 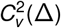 vs *s*_*b*_, adder strategy shows no-correlation and timer-like shows a negative correlation. To understand better the properties of the robust oscillations on the dynamics of the central moments presented in Figure 3.a and 3.b., we studied the auto-correlation fuction of the size. This auto-correlation *γ*(*t′*) is defined through the formula:

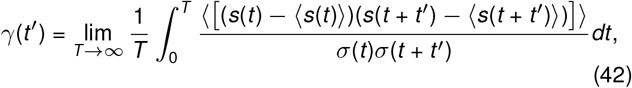

with *σ*(*t*) being the standard deviation of the size at time *t* and ⟨*x*⟩ is the mean value of the random variable *x*

**Figure 3:**
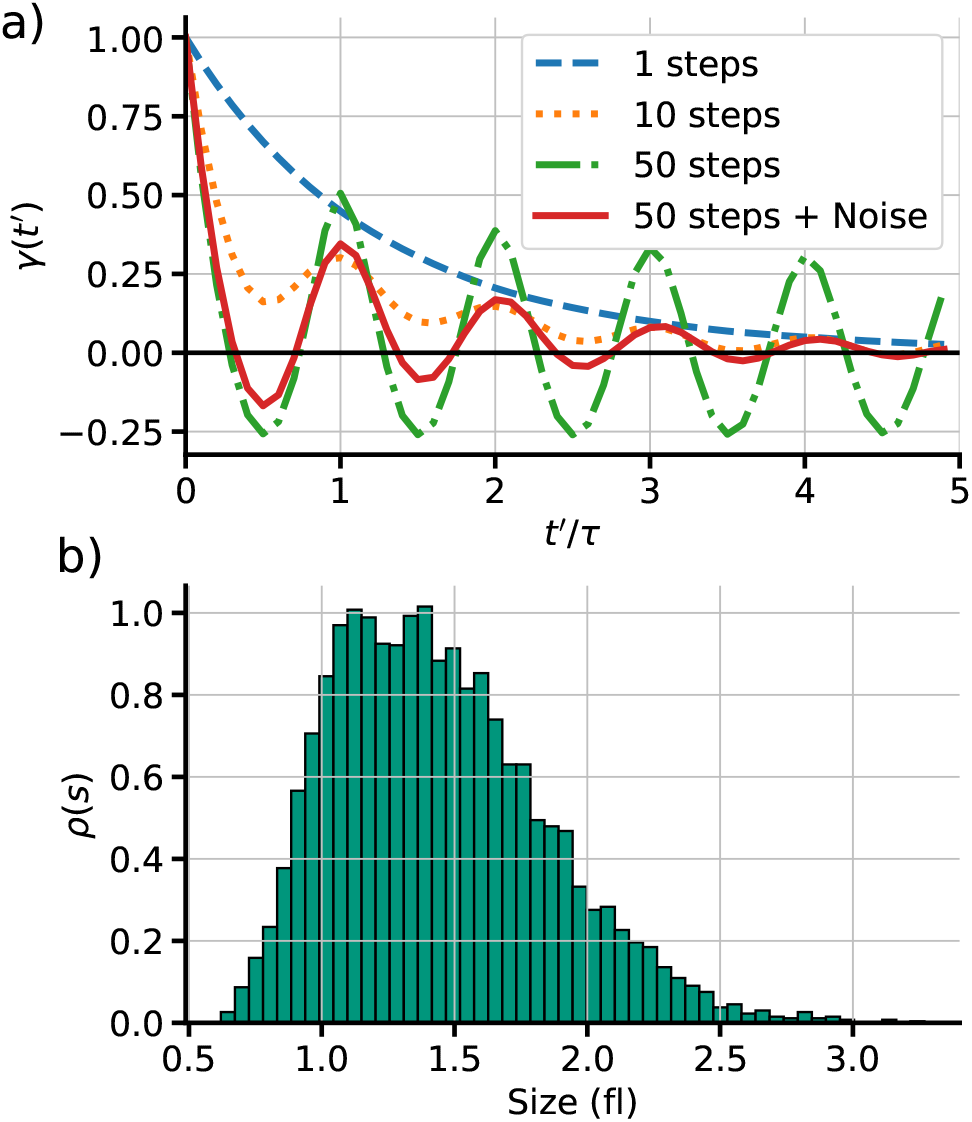
a) Cell size auto-correlation *γ*(*t′*) for different time periods *t′* for four different division conditions: one division step (dashed, blue line), 10 division steps (dotted, orange line), 50 division steps (green, dash-dot line) and 50 division steps including noise in growth rate and septum position (red continuous line). b) Simulation of stationary state of the histogram of bacterial size with all the noise sources considered.

How the auto-correlation dynamics changes along the time is presented on Figure 3. a. where we present four different cases: a single division step, which size dynamics have been presented already [22]. This auto-correlation decays exponentially to zero. By increasing the division steps, for instance to 10 steps, oscillations appear around a decaying trend. For a division almost deterministic, for instance 50 steps, these oscillations have higher amplitude around zero. When noise on both growth rate and septum position is considered, these oscillations are damped in the same way found in size dynamics, converging to zero. This asymptotic decorrelation let the distribution reach a stationary distribution (at *t* = 10*τ*) with fixed moments which is presented in Figure 3. b. for *M* = 10 and the noises explained above.

## Discussion

In this article, we present a theoretical scheme to estimate the stochastic dynamics of the cell size for a population of constant number. This approach, based on the Chapman-Kolmogorov formalism, assumed that the population number remains constant along the time. Although the framework can be extended to a exponentially growing population, simulation could be unstable since the number of cells in these populations grows exponentially on time. We think to consider this case of CKE in future studies.

Additionally to the estimation of size dynamics, our framework can be also used for: Simulate most of the division strategies found in *E. coli* : timer-like, adder and sizer-like [10, 32]. Estimate the distribution of division times, size at division and size at birth. Noise in both septal ring placing and cell-to-cell growth rate can be considered as well [26].

As is shown in Figure 2, oscillations in both, the mean size ⟨*s*⟩ and 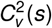, were found. When only stochasticity on division times is considered, these oscillations are maintained over an arbitrary long period of time having lower amplitude when an initial size distribution with finite variance is considered. Some damping occurs if other sources of noise like the cell-to-cell growth rate variability and septum position are added.

This robustness on the oscillations can be understood as result of the asymptotic periodic properties of the probability *P_n_* of have *n* divisions at time *n*. These probabilities are invariant under the simultaneous transformation *n* → *n* + 1 and *t* → *t* + *τ* originating the oscillations. This invariance under this transformation, is broken when other noise sources are considered.

These properties found in size dynamics can be also obtained using the classical Discrete Stochastic Maps. A clear correspondence between DSM and our model can be found when the adder strategy is considered. In this case, the stochastic map between size at birth *s_b_* and size at division is:

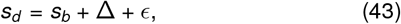

with *ϵ* being an independent random variable with zero mean and a distribution fitted from experiments. If exponential growth is considered, the cell-cycle duration *τ_d_*, as random variable, can be obtained from (43):

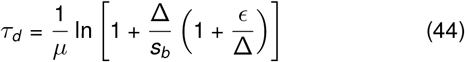

where, using (22), some analogy to (32) can be found. Using these times with parameters fitted from the data, similar oscillations in both size trends and auto-correlation can be obtained.

The main difference with DSMs is found where deviations from the adder are considered. Using the CRMs, the fluctuations (*ϵ*) in the division strategy (43) will depend on the size at birth unlike the DSM where these fluctuations are not related to any other variable. Some preliminary observations on the dependence of the fluctuations on added size with the size at birth in sizer-like division in *E coli* have been reported [14, 36] but further observations are needed.

Including cell size stochasticity to gene expression can be an important tool to understand the origin of the fluctuations in molecule concentration. Some efforts have already been done to understand these effects in simple regulatory networks [37–39] but our formalism can improve this study to more complex gene regulatory architectures. Other effects such as the division strategy, the noise in growth rate and the asymmetric cell splitting can be also studied using our framework.

## Appendix

**Figure A1:**
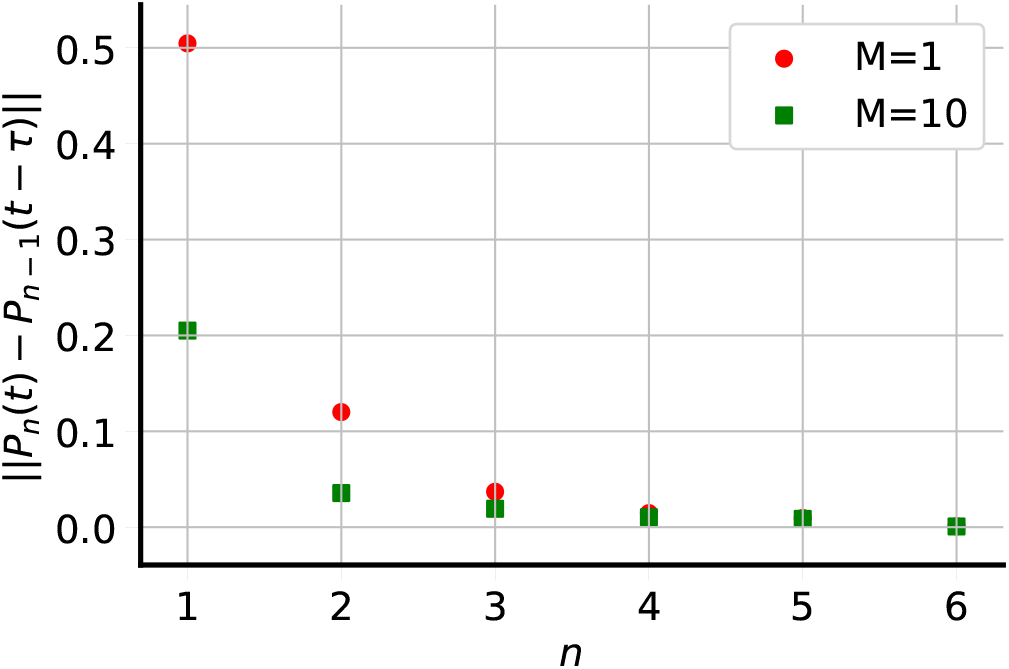
Distance between the probabilities *P_n_*(*t*) and *P*_*n*−1_(*t − τ*) for two different division steps (red dots: *M* = 1) (green squares: *M* = 10).

## A The asymptotic periodicity of the system

To check the property (41), we calculate numerically the distance between the probabilities *P_n_*(*t*) and *P*_*n−*1_(*t − τ*) using the expression:

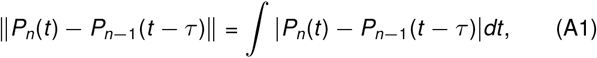

the results are presented in Figure A1 where we can check how this distance decays asymptotically to zero as *n* increases.

## B Size dynamics for different growth conditions and division rates

Past studies suggested that the periodicity under translations in *τ* and *n* is an exclusive property of the exponential growth [33]. Thus, if other growth conditions are considered, the symmetry under these translations is broken and the robust oscillations are no expected.

We simulated the size dynamics of different possible growth and division rates in similar way explained in past studies [17]. Thus, we define the growth law as (2), where exponential growth is defined as *g*(*s*, *t*) = *μs* and the linear growth is given by *g*(*s*, *t*) = *μ* with *μ* being a constant.

On the other hand, the division rate is defined by the function *h* as (6). In figure A2, we compare two division rates for the linear growth, one of these are *h* = *k* and the other is *h* = *ks* with k a constant. These rates define the occurrence of a given division steps. In figure A2 we considered *M* = 10 steps to trigger the division.

In Figure A2.a. we present the dynamics of the mean cell size ⟨*s*⟩ along the time with some single trajectories in the background (gray lines) presented to compare them to the mean trend. While in Figure A2.c. we present its variability 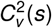 along the time for for a linear growth *g* = *μ* and constant division rate *h* = *k*. b) 〈*s*〉 vs *t* and d) 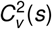 vs *t* both for a linear growth *g* = *μ* and a division rate porportional to the cell size *h* = *ks*. Perfectly symmetric splitting and no-noise in growth rate is considered.

**Figure A2:**
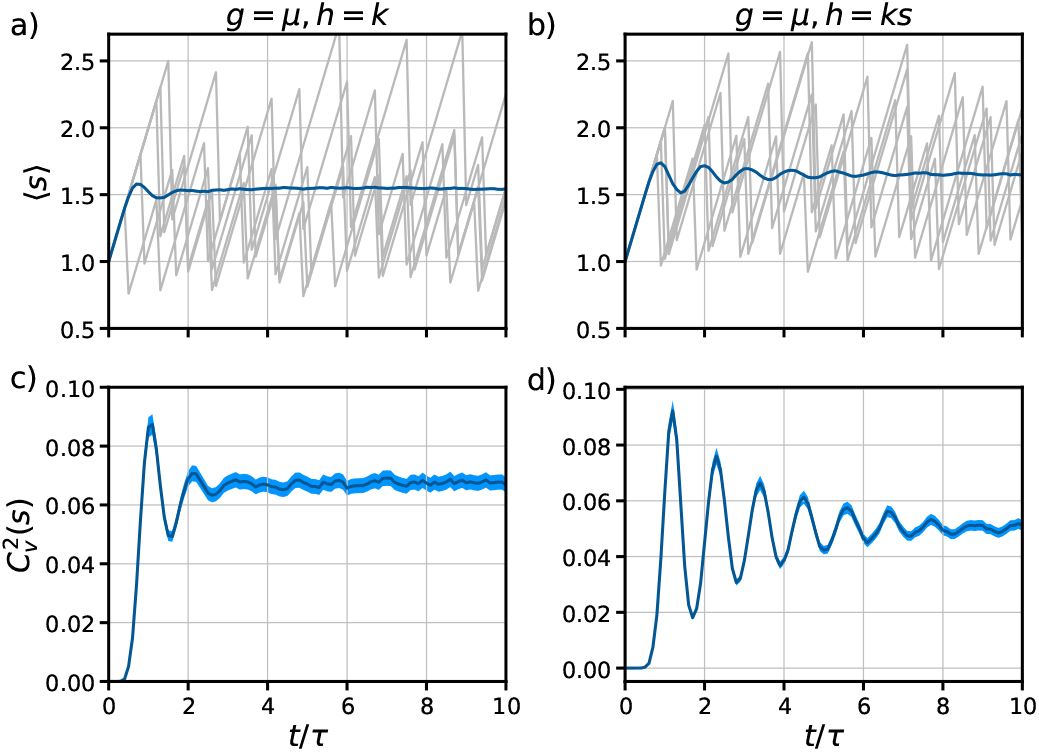
Mean cell size 〈*s*〉 and its variability 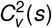 along the time for different growth and splitting rates. a) 〈*s*〉 vs *t* and c) 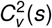 vs *t* both for a linear growth *g* = *μ* and constant division rate *h* = *k*. b) 〈*s*〉 vs *t* and d) 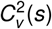 vs *t* both for a linear growth *g* = *μ* and a division rate proportional to the cell size *h* = *ks*.

